# Impaired perceptual learning in Fragile X syndrome is mediated by parvalbumin neuron dysfunction in V1 and is reversible

**DOI:** 10.1101/217414

**Authors:** Anubhuti Goel, Daniel A. Cantu, Janna Guilfoyle, Gunvant R. Chaudhari, Aditi Newadkar, Barbara Todisco, Diego de Alba, Nazim Kourdougli, Lauren M. Schmitt, Ernest Pedapati, Craig A. Erickson, Carlos Portera-Cailliau

## Abstract

Atypical sensory processing is a core characteristic in autism spectrum disorders^1^ that negatively impacts virtually all activities of daily living. Sensory symptoms are predictive of the subsequent appearance of impaired social behavior and other autistic traits^2, 3^. Thus, a better understanding of the changes in neural circuitry that disrupt perceptual learning in autism could shed light into the mechanistic basis and potential therapeutic avenues for a range of autistic symptoms^2^. Likewise, the lack of directly comparable behavioral paradigms in both humans and animal models currently limits the translational potential of discoveries in the latter. We adopted a symptom-to-circuit approach to uncover the circuit-level alterations in the *Fmr1^-/-^* mouse model of Fragile X syndrome (FXS) that underlie atypical visual discrimination in this disorder^4, 5^. Using a go/no-go task and in vivo 2-photon calcium imaging in primary visual cortex (V1), we find that impaired discrimination in *Fmr1^-/-^* mice correlates with marked deficits in orientation tuning of principal neurons, and a decrease in the activity of parvalbumin (PV) interneurons in V1. Restoring visually evoked activity in PV cells in *Fmr1^-/-^* mice with a chemogenetic (DREADD) strategy was sufficient to rescue their behavioral performance. Finally, we found that human subjects with FXS exhibit strikingly similar impairments in visual discrimination as *Fmr1^-/-^* mice. We conclude that manipulating orientation tuning in autism could improve visually guided behaviors that are critical for playing sports, driving or judging emotions.

---

For these studies, we focused on FXS because it is the leading inherited cause of autism^**5**^, because there are no associated major neuroanatomical defects, and because a single, well-characterized animal model, the *Fmr1^-/-^* mouse, is widely used. Although many human psychophysical studies have demonstrated deficits in visual perception in individuals with autism, including those with FXS^6, 7^, whether animal models of autism also exhibit impaired visual processing is not known. Thus, we first sought to determine whether *Fmr1^-/-^* mice manifest perceptual learning deficits associated with abnormal visual sensory discrimination. We trained male and female *Fmr1* knockout (*Fmr1^-/-^*; n= 21) and wild-type (WT; n= 19) mice (FVB strain) on a go/no-go visual discrimination task^8, 9^. Following water deprivation, awake head-restrained young adult mice (2-4 months old) were allowed to run on an air-suspended polystyrene ball while they performed the task (**Fig. 1a, b**; see **Materials and Methods**). Mice were presented with sinusoidal gratings drifting in two orthogonal directions, 45° (preferred, ‘go’) vs. 135° (non-preferred, ‘no-go’) at 100% contrast. Incorrect behavioral responses resulted in a 6.5 s ‘time-out’ period (**Fig.1c**). Task performance, as determined by the discriminability index statistic *d’* (**Materials and methods**), was dependent on primary visual cortex (V1), because pharmacological silencing of V1 with bilateral infusions of muscimol, a GABA-A receptor agonist, reversibly disrupted perceptual learning in WT mice (**Fig. 1d**).

**Figure 1:**
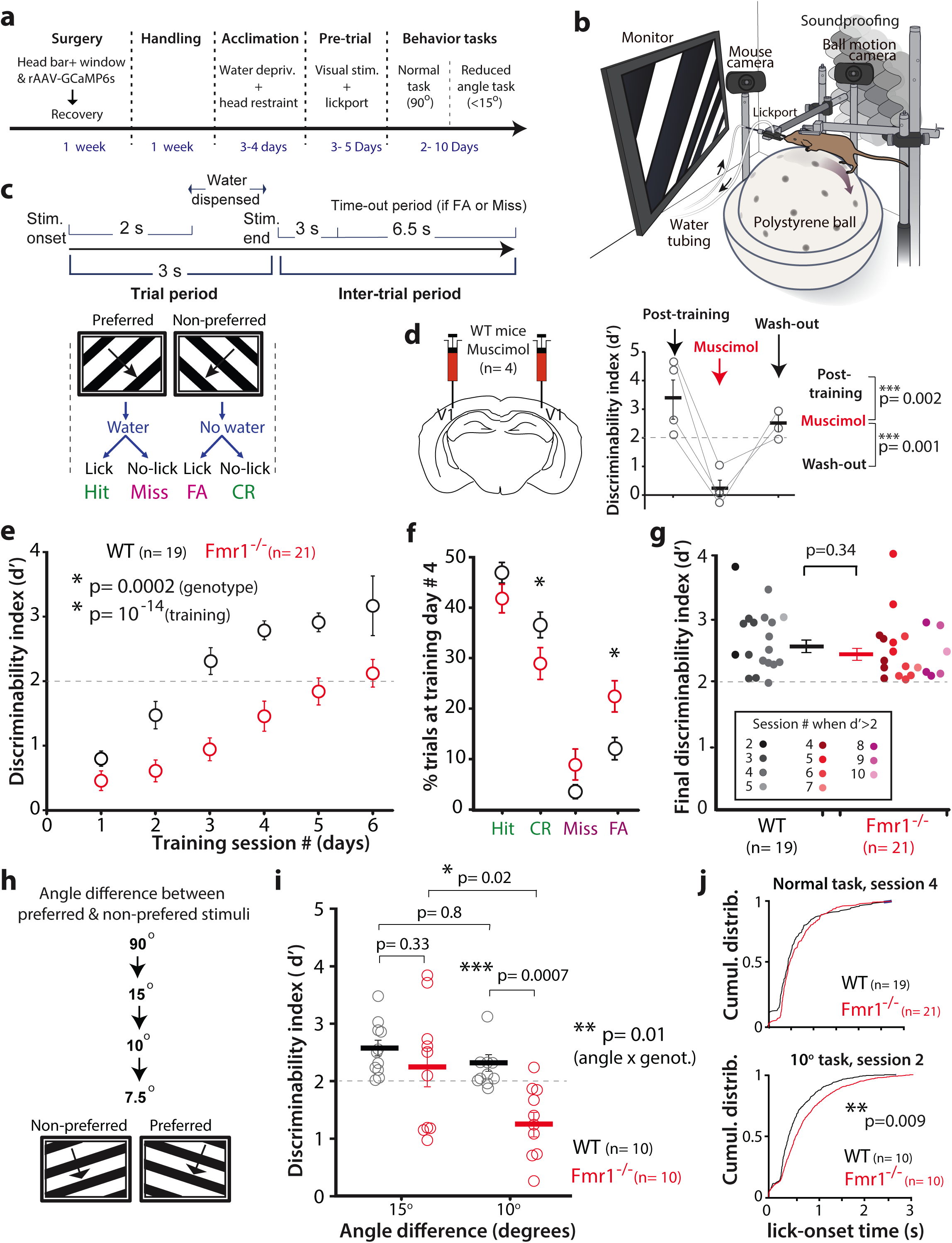
***Fmr1^-/-^ mice* have impaired performance in a motion visual discrimination task.** a. Experimental design and timeline for the visual discrimination task.
b. Cartoon of the behavioral apparatus.
c. Timeline of an individual trial for the go/no-go visual discrimination task in water deprived mice. FA: false alarm; CR: correct rejection.
d. Bilateral inactivation of V1 with muscimol impairs task performance in expert WT mice.
e. *Fmr1^-/-^* mice are delayed in learning of the basic task (90^o^ difference between preferred and non-preferred stimuli). The dashed line at d’= 2 indicates expert performance threshold (equivalent to ≥90% correct rejections). Error bars in this figure indicate the s.e.m. A repeated measures ANOVA test was used for panels e and i.
f. On the final test session at 90^o^ (prior to being advanced to reduced angle task), the discriminability index (d’) was equivalent between genotypes. In panels f & i, symbols represent different mice. Unpaired, one-tailed Student t-test was used for panels f-g;
g. Lower task performance of *Fmr1^-/-^* mice on day #4 is associated with a significantly higher proportion of false alarm responses and lower proportion of correct rejections.
h. A reduced angle task was implemented after mice learn the 90^o^ task.
i. Selective impairment of *Fmr1^-/-^* mice in task performance when the angle difference between preferred and non-preferred stimuli drops below 15^o^.
j. *Fmr1^-/-^* mice show a significant delay in lick onset time on the reduced angle task relative to WT mice.

WT mice learned quickly (3-4 sessions) to lick in response to the preferred orientation for a water reward and withhold licking when presented with the non-preferred orientation (**Fig. 1e**). In contrast, *Fmr1^-/-^* mice exhibited a significantly delayed learning curve, compared to the WT mice (**Fig. 1e** and **Suppl. Fig. 1**; F_1,29_= 1.76, p= 0.0002, two-way ANOVA with repeated measures on training). *Fmr1*^-/-^ mice exhibited a significantly higher percentage of false alarm (FA) responses compared to WT mice at session #4 (**Fig. 1f** and **Suppl. Fig. 2**; 11.3 ± 2.2% in WT vs. 22.3 ± 3.1% in *Fmr1*^-/-^; p= 0.009, Students t-test), which likely contributed to their poor performance during early training sessions. This increase in FA rates in *Fmr1^-/-^* mice was not caused by hyperactivity or abnormal locomotion, as the average running speed was similar between mice of both genotypes (not shown) and there was no change in running speed in mice of either genotype during the course of each trial on session 1 (**Suppl. Fig. 3c**). However, by session 4, we observed significantly more slowing down in WT than in *Fmr1^-/-^* mice, towards the end of each trial (**Suppl. Fig. 3c**). The delay of *Fmr1^-/-^* mice in learning the visual discrimination task was evident in female and male mice alike (**Suppl. Fig. 4**). Importantly, both WT and *Fmr1^-/-^* mice exhibited significant improvements in task performance throughout training (F_(4,116)_= 4.63, p< 10^-14^, two-way ANOVA with repeated measures on training session). Even though *Fmr1^-/-^* mice took, on average, 2.5 sessions longer to achieve a d’ > 2 (**Fig. 1e;** 3.5 ± 0.2 for WT vs. 6.0 ± 0.4 for *Fmr1^-/-^*; p= 4.3 x 10^-6^, t-test), there was no significant difference in the final d’ values between WT and *Fmr1^-/-^* mice (**Fig.1 g**). Thus, *Fmr1^-/-^* mice eventually achieve the same level of performance in this visual discrimination task as WT controls. Notably, when we reduced the contrast of gratings, *Fmr1*^-/-^ mice did not exhibit obvious impairments in visual perception, at least down to 10% contrast (**Suppl. Fig. 5**), suggesting that their delayed learning was not due to a primary visual deficit.

*Fmr1^-/-^* mice are known to exhibit a broadening of receptive fields in somatosensory cortex^10-12^. Similar broader tuning in V1, if it exists, could affect the discrimination of visual stimuli with very similar orientations. Therefore, we next tested whether *Fmr1^-/-^* mice would be particularly challenged by a reduced angle task, in which the difference in angle between the preferred and non-preferred orientation was gradually reduced to 7.5^o^, after the animals had learned the basic 90^o^ task (**Fig. 1h**). A difference in orientation angle of 15^o^ did not impair the performance of either WT or *Fmr1^-/-^* mice (n= 10 for each); however, a further reduction down to 10^o^ resulted in a significant reduction in *d’* values of *Fmr1^-/-^* mice, but not in WT controls (**Fig. 1i**; interaction effect, F_(1,18)_= 7.42, p= 0.01, ANOVA; d’ at 10^o^ was 2.3 ± 0.1 for WT vs. 1.2 ± 0.2 for Fmr1^-/-^; post-hoc t-test, p= 0.0007). A reduction in the angle difference to 7.5^o^ further impaired the performance of *Fmr1^-/-^* mice, but also led to a decrease in d’ in some of the WT mice (**Suppl. Fig. 6**). To further probe the extent to which mice were challenged by this reduced angle task, we assessed their response times and observed a significant delay in the distribution of licking onset in *Fmr1^-/-^* mice, compared to WT mice, for the reduced angle task, but not for the normal (90^o^) task (**Fig. 1j**, K-S test, p= 0.009). This suggests that *Fmr1^-/-^* mice take longer to make a decision only in the face of ambiguous sensory information.

Having established a defect in perceptual learning in the fragile X mouse model that is relevant to the human disease, we next adopted a reverse engineering approach to identify the circuit- and neuronal-level alterations that might underlie the impaired visual discrimination. In light of various reports of cortical hyperexcitability and network hypersynchrony in *Fmr1^-/-^* mice^13-15^, we first investigated whether the perceptual learning deficit we observed in *Fmr1*^-/-^ mice, was caused by abnormal orientation tuning of pyramidal cells in V1. To test this, we performed in vivo 2-photon calcium imaging in layer (L) 2/3 neurons in awake mice running on a floating polystyrene ball (**Fig. 2a-c**; **Materials and methods**). A rAAV to express GCaMP6s^16^ was injected in V1 following stereotaxic coordinates, and successful targeting was confirmed using intrinsic signal imaging (**Fig. 2b**). We recorded both spontaneous and visually evoked activity in L2/3 neurons (**Fig. 2c**). For the latter, WT and *Fmr1^-/-^* mice (n= 9 and 10, respectively) were presented with 4 sets of sinusoidal gratings drifting in 8 different directions, at random (**Fig. 2d**; **Materials and methods**). Although previous studies have reported hyperexcitable cortical circuits in *Fmr1*^-/-^ mice (reviewed in ^13^), we did not observe a significant increase in either spontaneous or visually evoked activity in *Fmr1*^-/-^ mice (**Fig. 2e** and **Suppl. Fig. 7a, b**). We did find a trend toward higher population coupling in *Fmr1*^-/-^ mice (**Suppl. Fig. 7c**), which is consistent with published results showing local circuit hypersynchrony in these mice^14^.

**Figure 2:**
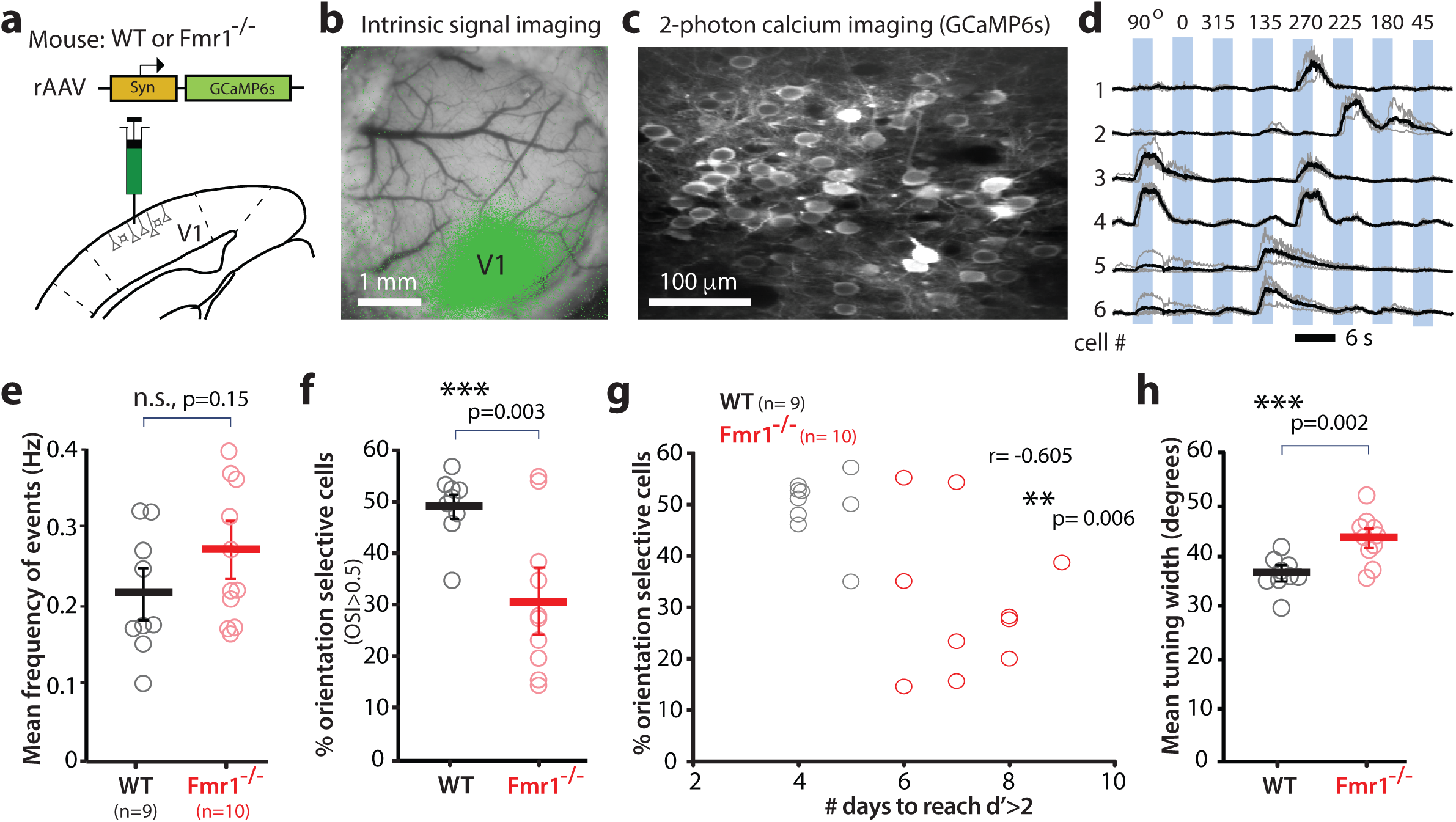
**Orientation tuning deficits in V1 correlate with task performance in *Fmr1^-/-^* mice.** a. Cartoon of rAAV-GCaMP6s injection into V1.
b. Intrinsic signal imaging was then performed 2-3 weeks after rAAV injection to confirm appropriate targeting of V1 (green map).
c. Representative field of view for in vivo two-photon calcium imaging experiment in V1. Imaging was performed 3-4 weeks after rAAV injection at 15 fps.
d. Example traces of changes in GCaMP6s fluorescence intensity (ΔF/F) for 6 representative neurons in V1 that exhibit a range of responses from narrow tuning (cell 1) to broad tuning (cells 2 & 3). Responses to single trials are shown in gray, averages of 4 responses are in black.
e. Visual evoked activity (as measured by the frequency of fluorescence peaks) is similar between WT and *Fmr1^-/-^* mice. Symbols in panels e-h represent different mice. Unpaired, one-tailed Student t-test was used for panels e-f, h.
f. The percentage of orientation selective neurons in V1 is significantly lower in *Fmr1^-/-^* mice.
g. Inverse correlation between the percentage of orientation selective neurons in V1 and performance on the visual discrimination task (as measured by the number of days required to reach a d’>2).
h. The mean orientation tuning width for V1 neurons is significantly higher in *Fmr1^-/-^* mice. Tuning width also correlates with task performance (see **Suppl. Fig. 8**)

Despite the seemingly normal frequency of visually evoked activity in *Fmr1*^-/-^ mice, mutant mice had a significantly lower percentage of orientation selective (OS) cells in L2/3 (**Fig. 2f**; 49.5 ± 2.1% in WT vs. 31.2 ± 4.6% in *Fmr1*^-/-^; p= 0.003, t-test). In other words, on average, pyramidal neurons in *Fmr1^-/-^* mice were tuned to multiple orientations. Importantly, when we trained these mice on the visual discrimination task, we found a significant inverse correlation between the percentage of OS cells and the number of days it took animals to reach a d’ > 2 (**Fig. 2g**; r= -0.605, p= 0.006). This implies that, with fewer available OS cells in V1, *Fmr1*^-/-^ mice had more difficulty discriminating between two different orientations, particularly when the difference was small (**Fig 1 h-j**). In addition, in vivo calcium imaging revealed that L2/3 neurons in V1 of *Fmr1^-/-^* mice had a significantly broader tuning compared to those in WT mice (**Fig. 2h;** 36.7 ± 1.0^o^ in WT vs. 43.3 ± 1.4^o^ in *Fmr1*^-/-^; p= 0.002, t-test). This 6.6^o^ difference in the mean tuning width of pyramidal neurons in V1 between WT and *Fmr1*^-/-^ mice, though slight, might be sufficient to explain why *Fmr1^-/-^* mice can discriminate at 15^o^ but not at 10^o^. Critically, we also found a significant correlation between the tuning width of L2/3 cells and the number of days it took the animals to reach a d’ > 2 (**Suppl. Fig. 8**; r= 0.48, p= 0.041).

Abnormal V1 network dynamics pertaining to orientation selectivity and tuning width could be the result of dysfunction in parvalbumin (PV) interneurons, the most prevalent inhibitory neuron in V1^17^. PV cells exhibit very broad orientation tuning by simply responding to all orientations, since they receive local input from a wide range of orientation tuned pyramidal cells^18-20^. Furthermore, selective stimulation of PV cells in V1 with channelrhodopsin-2 leads to improved feature selectivity and visual discrimination^9^. For these reasons, we tested the hypothesis that PV cells were hypoactive in fragile X mice. We used in vivo calcium imaging to record the activity of PV neurons in V1 of WT and *Fmr1*^-/-^ mice (n= 6 and 7, respectively) that expressed Td-Tomato in PV neurons (PV-Cre mice x ai9 mice; see **Materials and methods**). At the time of the cranial window surgery, we injected a Cre-dependent virus into V1, to selectively express GCaMP6s in PV cells (**Fig. 3a, b**). Our calcium imaging recordings revealed stark differences in the activity of PV cells between WT and *Fmr1^-/-^* mice; whereas traces of PV cell activity in WT mice showed the expected broadly tuned, non-selective responses to visual stimuli, traces of PV cells in *Fmr1*^-/-^ mice exhibited little visually evoked activity (**Fig. 3c**). While there was no significant difference in the amplitudes of calcium transients in PV cells in *Fmr1^-/-^* mice (**Fig. 3d**), we found a significantly lower frequency of events triggered by visual stimuli (**Fig. 3e**; 1.8 ± 0.5 for WT vs. 0.3 ± 0.1 for *Fmr1*^-/-^; p= 0.02, t-test). One of our criteria for selecting PV cells for analysis in both WT and *Fmr1*^-/-^ mice was that they exhibit at least one calcium transient in the recordings (**Materials and methods**), and neither the proportion of active PV cells (**Fig. 3f**; p= 0.5, t-test), nor the amplitude and frequency of spontaneous calcium transients in PV cells, were significantly different between WT and *Fmr1^-/-^* mice (**Suppl. Fig. 9**). We also found a significantly lower fraction of stimulus-responsive PV cells in *Fmr1*^-/-^ mice (**Fig. 3g**; 0.7 ± 0.02 for WT vs. 0.4 ± 0.03 for *Fmr1*^-/-^; p < 10^-5^, t-test), which would also ultimately be expected to affect the functional output of V1.

**Figure 3:**
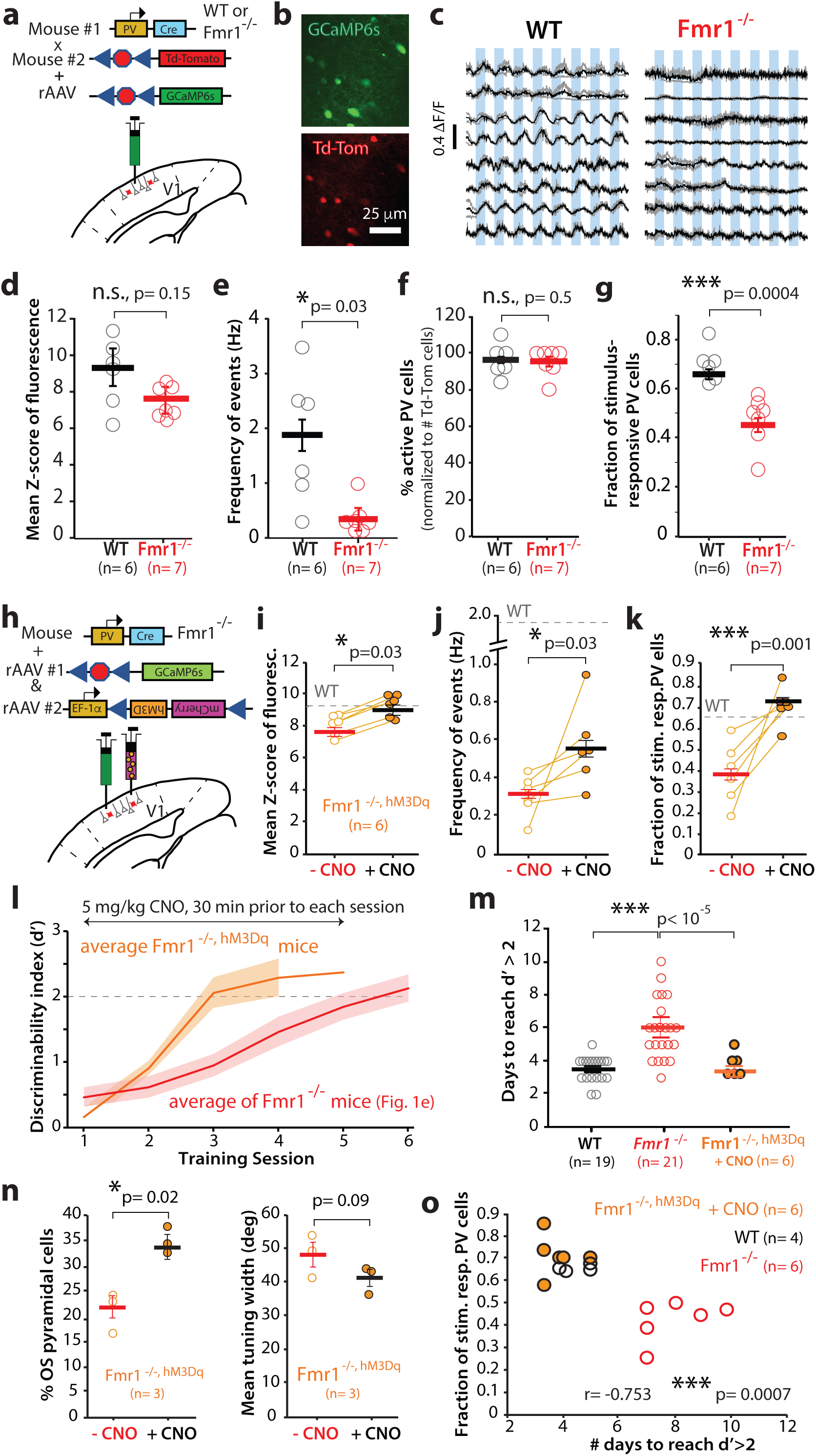
**A DREADD strategy to correct the hypoactivity of parvalbumin interneurons in V1 rescues task performance in *Fmr1^-/-^* mice.** a. Cartoon of strategy for selective GCaMP6s expression in PV interneurons.
b. Representative field of view for in vivo 2-photon calcium imaging in PV neurons expressing GCaMP6s (green) and Td-Tom (red).
c. Example traces of changes in GCaMP6s fluorescence intensity (ΔF/F) for 8 representative PV neurons in V1 from 4 WT (left) and 4 *Fmr1^-/-^* mice (right). Responses to 8 different directions from single trials are shown in gray, while the averages of 4 trials are in black.
d. Visual evoked activity (as measured by the mean fluorescence Z-scores) is similar between WT and *Fmr1^-/-^* mice (sample size in parenthesis). Symbols in panels d-j represent different mice. Unpaired, one-tailed Student t-test was used for panels d-g.
e. The frequency of visually evoked calcium transients in PV neurons is significantly lower in *Fmr1^-/-^ mice.*
f. WT and *Fmr1^-/-^* mice and similar percentages of PV cells that were active.
g. The fraction of visually-responsive PV cells is significantly reduced in *Fmr1^-/-^* mice. There is an inverse correlation between the fraction of stimulus responsive PV cells and behavioral performance (panel o).
h. Cartoon of strategy for selective rAAV-EF1a-DIO-hM3D(Gq)-mCherry expression in PV interneurons of *Fmr1^-/-^* mice.
i, j. The activity of PV cells (as measured by both median fluorescence Z-score, *j*, or the frequency of calcium transients, in *Fmr1^-/-, HM3Dq^* mice increases significantly after CNO administration. Symbols in panels j-l represent different mice. A one-tailed, unpaired Student t-test was used for panels i-k.
k. The fraction of stimulus-responsive PV cells also increases significantly after CNO. Note that the fraction of visually responsive PV cells was comparable between *Fmr1*^-/-^ mice expressing DREADDs (before CNO) and *Fmr1*^-/-^ mice in panel g (also see **Suppl. Fig. 10 e**)
l. *Fmr1^-/-, HM3Dq^* mice (n= 6; treated with CNO 30 min prior to each session) learned the basic 90^o^ task in ~3 d on average. The rate of learning for *Fmr1^-/-^* mice (from Fig. 1e) is shown for comparison. The solid line indicates the mean, and the shaded area shows the standard error. The dashed line at d’= 2 indicates expert performance threshold.
m. The percentage of OS pyramidal neurons in *Fmr1^-/-, HM3Dq^* mice was significantly higher after CNO administration, but there was no obvious effect on their tuning width. Student t-test.
n. *Fmr1^-/-, HM3Dq^* mice treated with CNO learned the basic 90^o^ task significantly faster than *Fmr1^-/-^* mice and as fast as WT mice. Repeated measures ANOVA.
o. There is a strong inverse correlation between task performance (days to reach d’>2) and the fraction of stimulus-responsive PV cells in V1.

Based on the finding that PV cells were indeed hypoactive in *Fmr1^-/-^* mice, we hypothesized that a successful manipulation of PV cell activity that would restore their output in these animals, might also improve their performance on the visual discrimination task. Hence, we used a Designer Receptors Exclusively Activated by Designer Drugs (DREADD)^21^ approach (see **Materials and methods**) to selectively express the excitatory hM3Dq receptor in PV cells of *Fmr1^-/-^* mice (n= 6; **Fig. 3h**). We then used the hM3Dq ligand, clozapine-N-oxide (CNO, 5 mg/kg, i.p.), to excite PV cells and increase their output in these *Fmr1^-/-, hM3Dq^* mice. Overexpressing hM3Dq in PV cells alone (before administering CNO) did not affect visually evoked activity of PV cells in *Fmr1^-/-, hM3Dq^* mice (**Suppl. Fig. 10a-d**). In contrast, 30 min after a single CNO injection, we observed a robust increase in visually evoked PV cell output in these *Fmr1*^-/-, hM3Dq^ mice (**Fig. 3i-k**, **Suppl. Fig. 10e**). Specifically, we observed a significant increase in both the z-score of the amplitude of visually evoked calcium transients in PV cells of *Fmr1*^-/-, hM3Dq^ mice (**Fig. 3i**; 7.9 ± 0.3 before CNO vs. 8.9 ± 0.3 after CNO; p= 0.03, t-test), and in the frequency of those events (**Fig. 3j**; 0.3 ± 0.04 before CNO vs. 0.6 ± 0.1 after CNO; p= 0.03, t-test). The fraction of stimulus responsive PV cells in *Fmr1*^-/-, hM3Dq^ mice was also significantly increased by CNO, restoring it to WT levels (**Fig. 3k**; 0.4 ± 0.06 before CNO vs. 0.7 ± 0.04 after CNO; p= 0.001, t-test). Importantly, the fact that we could increase the activity of PV cells with DREADDs supports the notion that PV cells were not silent in *Fmr1^-/-^* mice due to poor health. Also, the proportion of PV cells that was active did not change after CNO administration (not shown), suggesting that the DREADD effect on the fraction of visually responsive PV neurons was not due to simply making previously silent cells more active.

Having restored visually evoked PV cell activity in *Fmr1*^-/-/, hM3Dq^ mice to near normal WT levels, we hypothesized that we might be able to reverse the delay in learning the visual discrimination task. A subset of the DREADD-expressing *Fmr1*^-/-^ mice were therefore trained on the standard visual discrimination task (90^o^ angle) and injected with CNO, ~30 min prior to each training session. This chemogenetic manipulation resulted in a leftward shift in the learning curve (i.e., faster learning) of CNO-treated *Fmr1*^-/-, hM3Dq^ mice (**Fig. 3l**), indicating that we were able to rescue the learning impairment by acutely elevating the PV cell output. CNO led to a significant reduction in the number of days required to reach expert level (d’ > 2) on the visual discrimination task compared to *Fmr1^-/-^* mice (**Fig. 3m**; 6.0 ± 0.4 d for *Fmr1^-/-^* vs. 3.7 ± 0.3 d *Fmr1^-/-, hM3Dq^* with CNO; F_2,43_= 1.7, p< 10^-5^, one-way ANOVA), which was comparable to the WT mice (3.5 ± 0.2; p= 0.53). To come full circle back to OS cells in V1, we also tested whether the DREADD manipulation on PV cells would be sufficient to affect the properties of pyramidal neurons in the circuit. Calcium imaging with rAAV-GCaMP6s in a group of *Fmr1^-/-, hM3Dq^* mice revealed that CNO administration significantly raised the proportion of orientation selective pyramidal cells and showed a trend towards sharper tuning (**Fig. 3n**; p= 0.03 and 0.07, respectively; t-test). Notably, the relationship between PV cell output and behavior was apparent from the negative correlation between the fraction of stimulus responsive PV cells and the number of days needed to reach a d’ > 2 (**Fig. 3o**; r -0.753, p= 0.0007). This relationship showed clearly how *Fmr1^-/-, hM3Dq^* mice treated with CNO were not distinguishable from WT mice.

It was recently argued that the absence of directly comparable behavior paradigms between human and animal studies is a real impediment to progress in translational research for autism^2^. It might even explain, in part, the failure of clinical trials in FXS^22^. In order to assess the translational potential of our findings of impaired visual discrimination (and, by extension, the associated circuit dysfunction) in a mouse model of FXS, we next asked whether the same perceptual learning task could be applied to humans with FXS. We implemented the same paradigm as in mice with relatively minor modifications, to make it suitable for individuals with FXS (**Fig. 4a, b**; **Materials and methods**). Healthy control human participants and FXS participants (n= 8 each; see **Suppl. Table 1**) were administered the task. Healthy controls learned the basic 90^o^ task with high discriminability very quickly (within the first ten trials) in a single training session (**Fig. 4c**). Performance declined slightly in some healthy control participants at reduced angles, but on average, this was not significant. In contrast, FXS participants showed significantly lower d’ at the 15^o^ task compared with 90 ^o^ or 45 ^o^ tasks (p= 0.002; repeated measures ANOVA). Thus, FXS participants and *Fmr1^-/-^* mice exhibit strikingly similar visual perception deficits for ambiguous stimuli with similar orientations. This suggests that a discrimination task like the one we used could eventually be used as biologically-based measure of sensory processing in human clinical trials.

**Figure 4:**
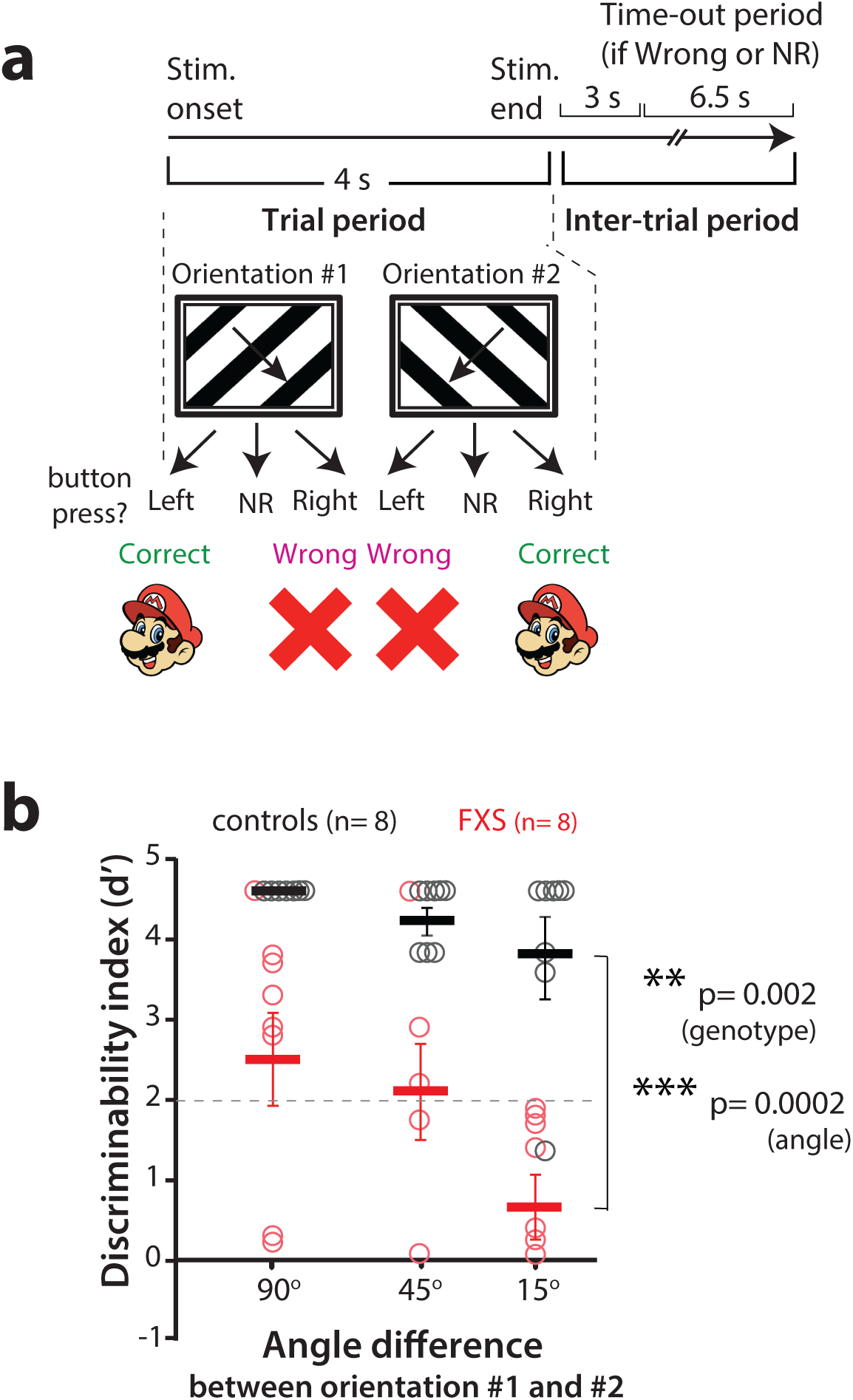
**Fragile X patients exhibit similar defects in visual discrimination as *Fmr1^-/-^* mice** a. Photograph of a FXS subject performing the visual discrimination task.
b. Timeline of an individual trial for the visual discrimination task in human subjects. NR: no response.
c. Task performance at different angles between orientation #1 and orientation #2 for FXS subjects and age-matched control participants. Individuals with FXS are able to perform the 90^o^ visual discrimination task with d’> 2 but they exhibit a significantly lower d’ than controls with the reduced angle task.

Progress in autism research is limited by the lack of clearly identified circuit-level alterations that can explain the neuropsychiatric phenotype that characterizes the disorder. Though circuit activity in monogenetic murine models of ASD can be readily interrogated and manipulated, there is increasing interest to demonstrate both face validity and predictive validity for these translational approaches to be used in clinical trials^2^. To bridge this gap, we implemented a fully translatable behavioral assay of sensory processing in both *Fmr1^-/-^* mice and FXS patients, and followed a symptom-to-circuit approach to delineate specific circuit-level defects using calcium imaging in V1. Our discovery that *Fmr1^-/-^* mice have a reduced proportion of orientation selective neurons with abnormally broad tuning in V1, as well as extremely hypoactive PV cells, provides a mechanistic understanding of their visual discrimination deficits. The fact that we could rescue their perceptual deficits in mice by restoring activity in PV cells with DREADDs and that humans with FXS exhibit analogous deficits in visual discrimination, provides a realistic path for novel translational clinical trials.

Our data implicates a role for PV cells in circuit dysfunction in FXS through converging evidence of their hypoactivity from experiments in two different groups of mice (*Fmr1^-/-^* and *Fmr1^-/-, hM3Dq^*), and from the DREADD approach, which not only restored PV cell activity, but also raised the percentage of orientation selective pyramidal cells in V1. It is exciting to consider that PV cell dysfunction could be involved in other aspects of FXS, such as impaired neuronal adaptation in tactile defensiveness^23^. Additionally, by rapidly translating this paradigm into a clinical population, we have substantially reduced potential barriers to further study whether PV cell dysfunction represents an important aspect (or even the principal one) of a *canonical micro-circuitry* in autism^2^.

## METHODS SUMMARY

Head-fixed, water-deprived, adult male and female WT and *Fmr1^-/-^* mice (2-4 months; FVB strain) were trained on a go/no-no task wherein they learned to discriminate between sinusoidal gratings drifting at two different orientations, 90^o^ apart. After learning the basic task, mice were advanced to a reduced angle task, in which the angle separating the orientations of preferred and non-preferred stimuli was reduced to 15^o^, 10 ^o^ and 7.5 ^o^. A subset of these mice received an injection of rAAV-GCaMP6s and a cranial window surgery, and then underwent two-photon calcium imaging at 15 Hz using a custom resonant scanning microscope. For PV cell imaging, we used PV-Cre x ai9 mice (Td-Tomato) and injected rAAV-syn-GCaMP6s or rAAV-fl-STOP-fl-GCaMP6s into V1. Analysis of calcium signals was performed with custom routines written in MATLAB. For DREADD experiments, we injected rAAV-EF1a-DIO-hM3D(Gq)-mCherry into PV-Cre x *Fmr1^-/-^* mice at the time of the cranial window surgery. We also trained these mice on the basic visual discrimination task. For human studies, we administered an analogous discrimination task to adolescent and adult participants with FXS and to age-matched healthy controls.

See below for a full description of methods.

## EXTENDED MATERIALS AND METHODS

### Experimental animals

All experiments followed the U.S. National Institutes of Health guidelines for animal research, under an animal use protocol (ARC #2007-035) approved by the Chancellor’s Animal Research Committee and Office for Animal Research Oversight at the University of California, Los Angeles. Experiments in Figs. 1 and 2 used male and female FVB.129P2 WT mice (JAX line 004828) and *Fmr1*^-/-^ mice^24^ (JAX line 004624) and experiments in Fig. 3 used male and female PV-Cre mice (JAX line 008069) that were crossed to the Ai9 (Td-Tom) reporter line (JAX line 007909) and the resulting PV-Cre x Ai9 mice were back crossed to FVB WT and *Fmr1*^-/-^ mice for 8 generations. All mice were housed in a vivarium with a 12/12 h light/dark cycle and experiments were performed during the light cycle. The FVB background was chosen because of its robust breeding, because FVB *Fmr1^-/-^* dams are less prone to cannibalizing their pups, and because FVB *Fmr1^-/-^* mice have well-documented deficits in sensory processing^13^. Additionally, to improve the survival of *Fmr1^-/-^* pups due to the possibility of littermates with different genotypes receiving unequal attention from the dam^25^ we used homozygous litters.

### Viral constructs

AAV1.Syn.GCaMP6s.WPRE.SV40 and AAV1.Syn.Flex.GCaMP6s.WPRE.SV40 were purchased from the University of Pennsylvania Vector Core and diluted to a working titer of 2e^13^ or 2e^12^ (to enable a longer period of optimal expression) with 1% filtered Fast Green FCF dye (Fisher Scientific). GCaMP6s was chosen over GCaMP6f because it detects more active neurons and it has an improved signal-to-noise ratio for more reliable spike detection^16^. For DREADD experiments, pAAV.hSyn.DIO.hM3D(Gq).mCherry was purchased from Addgene and diluted to a working titer of 2e^12^ with 1% Fast Green FCF dye.

### Cranial window surgery

Experiments were started with craniotomies performed at 6-8 weeks on the four different mouse lines mentioned above. Mice were anesthetized with isoflurane (5% induction, 1.5-2% maintenance via a nose cone) and placed in a stereotaxic frame. A 4.5 mm diameter craniotomy was performed over the right primary visual cortex (V1) and covered with a 5 mm glass coverslip, as previously described ^**26, 27**^. Before securing the cranial window with a coverslip, we injected ~50 nl of AAV1.Syn.GCaMP6s.WPRE.SV40 (Fig. 2, in vivo calcium imaging of L2/3 pyramidal neurons), or a cocktail of AAV1.Syn.GCaMP6s.WPRE.SV40 and AAV1.Syn.Flex.GCaMP6s.WPRE.SV40 (Fig 3, *in vivo* calcium imaging in PV cells), or a cocktail of AAV1.Syn.Flex.GCaMP6s.WPRE.SV40 and pAAV.hSyn.DIO.hM3D(Gq).mCherry (Fig. 3, to activate PV cells with DREADDs). Injections were performed ~ 3.0 mm lateral of lambda to target binocular V1. A custom U-shaped aluminum bar was attached to the skull with dental cement to head restrain the animal during behavior and calcium imaging.

### Optical intrinsic signal (OIS) imaging

Two weeks after cranial window surgery, OIS imaging was used to map the location of V1. Visual stimulation was provided by a piezo-actuator (Physik Instrumente) that deflected light from a red light emitting diode in front of the contralateral eye. The response for 30 stimulation trials was averaged, each consisting of 100 Hz deflections for 1.5 s. The response signal divided by the averaged baseline signal, summed for all trials, was used to generate the visual cortical map.

### *In vivo* two-photon calcium imaging

Calcium imaging was performed on a custom-built 2-photon microscope, with a Chameleon Ultra II Ti:sapphire laser (Coherent), resonant scanning mirrors (Cambridge Technologies), a 25X objective (1.05 NA, Olympus) and ScanImage software^28^. Mice were head restrained with a metal bar and allowed to run freely on a floating 8 cm polystyrene ball. Visual stimuli were generated using custom-written MATLAB (Mathworks) routines using Psychtoolbox that consisted of full-field square wave drifting gratings (2 cycles/s, 0.005 spatial frequency, 32 random repeats of 8 orientations) presented for 3 s and separated by a 3 s-long grey screen. Both spontaneous and visually evoked responses of L2/3 pyramidal cells from V1 were recorded at 15 Hz in 2-4 fields of view. Each FOV consisted of a median of 63 pyramidal cells (range 58-81; no differences between WT and *Fmr1^-/-^* mice) and/or 9 PV cells (range 8-12) in WT mice and 5 PV cells (range 3-8) in Fmr1^-/-^ mice. In each animal, imaging was performed at 2-3 depths (150-250 µm), and data was averaged from movies collected across all FOVs.

### Data analysis for calcium imaging

Calcium-imaging data were analyzed using custom-written MATLAB routines, which included modifications of our previously described MATLAB code^14^. X-Y drift in the movies was corrected using an iterative, cross-correlation-based, non-rigid alignment algorithm^29^. A semi-automated algorithm^16^ was used to select regions of interest (ROIs), each representing a single cell body, and to extract the fluorescence signal (*F*) for each ROI. A “modified Z-score” Z_F vector for each neuron was calculated as:

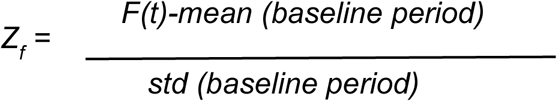

where the baseline period is the 10 s period with the lowest variation (standard deviation) in Δ*F*/*F*^23^. All subsequent analyses were performed using the Z_F vectors. Peaks were then detected in the Z-scores using the PeakFinder MATLAB script. These peaks were used to calculate the mean Z-score fluorescence (an estimate of amplitude of the fluorescence signal) and the frequency of events. Orientation selective cells were defined by an orientation selectivity index (OSI) calculated as:

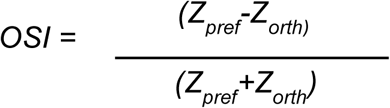

where Z_orth_ is the mean response to the orientation orthogonal to the preferred one (Z_pref_)^30^. A cell was considered orientation-selective if it had an OSI ≥ 0.5.

In order to determine whether an individual cell showed a time-locked or stimulus selective response to a visual stimulus in Fig. 3 and Suppl. Fig. 9c (which examines the correlation between the stimulus and the fluorescence signal in PV cells), we used a probabilistic bootstrapping method as described previously^23^. First, we calculated the correlation between the stimulus time-course and the Z_F vector, followed by correlation calculations between the stimulus time-course and 1,000 scrambles of all calcium activity epochs in Z_F (epoch = consecutive frames wherein Z_F ≥ 3). The 1,000 comparisons generated a distribution of correlations (R values), within which the correlation of the unscrambled data and the stimulus fell at a certain percentile. If the calculated percentile for a cell was less than 0.01, then we described that cell as being stimulus selective.

Correlations in Figures 2g, 3o and Sup. Fig. 8 were calculated using a Pearson correlation.

Population coupling (Suppl. Fig. 7c) was calculated as previously described^31^. It measures how individual neurons are correlated to the entire population of neurons. First, for each movie, we calculated a normalization factor to normalize for the different number of ROIs and activity levels within the movie. For the normalization procedure, the traces were first binned into 1 s time bins. If an ROI was active at least once (a peak detected) during this 1 s bin, it was converted to a ‘1’ in the binned trace. Then, we binned all these peak times into 1 s bins of binary values (0 or 1). These normalized traces were then used to calculate a Pearson correlation between each neuron and the entire population.

### Go/No-go visual discrimination behavioral paradigm in head-restrained mice

Awake, head-restrained young adult mice (2-4 months) were allowed to run on an air-suspended polystyrene ball while they performed the visual discrimination paradigm. Prior to performing the discrimination task the animals were subjected to handling, habituation and pretrial (Fig. 1a). After recovery from headbar/cranial window surgery, mice were handled gently for 5 min every day, until they were comfortable with the experimenter and would willingly transfer from one hand to the other to eat sunflower seeds. This was followed by water deprivation (giving mice a rationed supply of water once per day) and habituation to the behavior rig. During habituation, mice were head-restrained and allowed to run freely on the polystyrene ball in an enclosed sound proof chamber (**Fig. 1b**). Eventually, mice were introduced to the visual stimuli on the screen and the lickport (either commercial from Island Motion or custom-built at the UCLA electronics shop) that dispensed water (3-4 µL). This was repeated for 15 min per session for 2-3 days. Starting water deprivation prior to pretrials motivated the mice to lick (Guo et al., 2014). After habituation and ~15% weight loss, mice started the pretrial phase of the training. During pretrials, drifting sinusoidal gratings at 8 different directions were displayed on the screen. The monitor was placed at a distance of 20 cm from the mouse and stimuli were presented at random for 3 s, and each stimulus was coupled with a water reward dispensed through a lickport at 2 s after stimulus presentation. The mice were required to learn to associate a water reward soon after the stimulus was presented on the screen and that there was no water reward in the inter-trial interval (3 s period between trials). Initially, during pre-trials the experimenter pipetted small drops of water onto to the lickport to coax the mice to lick. Once the mice learned this and licked with 85% efficiency, they were advanced to the go/no-go task.

During the go/no-go visual discrimination task, drifting sinusoidal gratings (temporal frequency of 2 Hz, spatial frequency of 0.01 cycles /degree and 100% contrast) were displayed for 3 s, with water reward occurring 2 s after stimulus onset. Sinusoidal gratings of 45° and 135° orientations were randomly presented on the screen, but only the 45° orientation was coupled with a water reward (**Fig 1c**). Mice learned to discriminate between the two orientations and to lick in anticipation of the water reward during the 45° presentation (i.e., ‘go’) while withholding licking during the 135° orientation (i.e., ‘no-go’). Licking was recorded during the entire 3 s period; however, only licking between 2 s and 3 s contributed to the calculation of the behavioral response. Depending on the stimulus presented, the animal’s behavioral response was characterized as “Hit”, “Miss”, “Correct Rejection” (CR) or “False Alarm” (FA) (**Fig. 1c**). An incorrect response resulted in a time-out period of 6.5s, where nothing is presented on the screen. Training sessions consisted of 350 trials. To examine the psychometric threshold of task performance was tested at different contrasts (Sup. Fig. 5). d’ (discriminability index) was calculated as

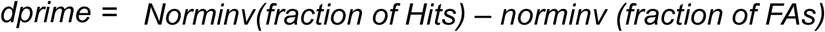

Custom-written routines and Psychtoolbox in MATLAB were used to present the visual stimuli, to trigger the lickport to dispense water, and to acquire data.

### Mouse locomotion video acquisition and analysis

Evenly distributed dots (1 cm in diameter) were painted on the surface of the polystyrene ball (**Fig. 1b**) to capture its motion with a high-speed webcam (Logitech C920). Custom MATLAB routines were used to analyze the ball motion videos. The webcam acquired 720 x 1280 videos at 60 fps. We ran each frame through a Hough transform to identify dark circles in the image (**Suppl. Fig. 3**). A series of spatial filters excluded irrelevant circles that were outside the ball’s circumference. A cap of 200 cm/s was set as a threshold above which there was a significant motion blur that led to inconsistent speed calculations (**Suppl. Fig. 3**). The algorithm for calculating ball motion speed first identified all the dots on the ball visible in each frame of the video, using spatial filters to isolate circular dots only on the ball. Then, it compared consecutive frames to determine how much each dot had moved relative to its position in the previous frame. Finally, the algorithm calculates the distance that dots moved and the median of overall dot movement over a 0.1 s interval to output absolute velocity of the ball. A separate script then aligned the velocities to the different stimulus presentations, which allowed us to compare mouse speeds at different points during the trial.

### Bilateral cannula implantation surgery

Similar to the cranial window surgery, adult WT mice were anesthetized with isoflurane and secured to a stereotaxic frame. We placed the mice on a heating blanket and used artificial tears to keep their eyes moist throughout the surgery. After exposing the skull, we drilled two shallow burr holes targeting V1 on both hemispheres (-4.0 mm posterior and +/- 3 mm lateral to Bregma). We then inserted 26 G guide cannulas (Plastics One) into the holes to a depth of -0.4 mm below the dura and the internal cannula projected further to -0.6 mm. After securing the cannula using cyanoacrylate glue (Krazy Glue) that was accelerated with cyanoacrylate glue accelerator (Insta-Set), dummy cannulas were screwed on to ensure that the guide cannula did not get clogged. Finally, we attached a headbar that would be used to immobilize the animal’s head during the visual behavior task. The cannula pedestals and headbars were further secured using dental cement that covered all exposed parts of the skull. After recovery, mice completed the visual discrimination task. Once the mice reached a d’ > 2, 8 µL muscimol solution (1 mg/ml in water) was infused bilaterally in isoflurane anesthetized mice through a 28 G internal cannula. After full recovery from anesthesia (<15 min), mice were immediately re-tested on the visual discrimination task. Two mice were also re-tested the following day to evaluate task performance after washout of muscimol. To confirm the correct targeting of the cannulas and muscimol infusions, mice were also infused by the fluorescent dye DiI, followed by transcardial perfusion with paraformaldehyde and histology (see below).

### Transcardial perfusion, DiI histology and PV immunohistochemistry

Mice were deeply anesthetized with isoflurane and underwent transcardial perfusion with 4% paraformaldehyde solution in sodium phosphate buffer (composition in mM: 30 mM NaH_2_PO_4_, 120 mM Na_2_HPO_4_, pH 7.4). Brains were kept overnight at 4°C first in 4% paraformaldehyde solution (pH 7.4) and then stored in 30% sucrose (in phosphate buffered saline at 4°C until sectioned.

### Human subjects

Eight males with FXS and eight male healthy controls, matched on chronological age, completed the visual discrimination experiment (**Supplemental Table 1**). Testing was conducted at Cincinnati Children’s Hospital Medical Center where the participants with FXS were originally recruited as part of our Center for Collaborative Research in Fragile X (U54). All FXS participants had full FMR1 mutations (>200 CGG repeats) confirmed by genetic testing. No participants had a history of non-febrile seizures or treatment with an anticonvulsant medication (see **Supplemental Table 2** for full medication details). FXS participants completed the Abbreviated Battery of the Stanford-Binet Intelligence Scales-Fifth Edition (SB-5). Control participants were recruited through community advertisements and were excluded for a history of developmental or learning disorders or significant psychiatric disorder (e.g., schizophrenia) in themselves or first degree-relatives, or for a family history of ASD in first- or second-degree relatives based on a brief screening interview. All participants >18 years of age provided written consent and minors provided assent, as well as written consent from their legal guardians. All study procedures were approved by the local Institutional Review Board.

Human FXS and control participants completed a visual discrimination task that was nearly identical to that used with mice with relatively minor modifications. Due to the additional cognitive demands of a go/no-go paradigm, including inhibitory control, which is known to be impaired in FXS^32^, we designed a forced two-choice visual discrimination task, so that all FXS participants could perform the task in a single session. It is conceivable however that FXS subjects could have learned the go/no-go task with subsequent training sessions, just as the mice required consecutive sessions to learn. Visual gratings were displayed on a 13-inch Hewlett Packard laptop computer with a 15-inch liquid crystal display and made responses on designated keys on the laptop keyboard. During the task, when the visual grating appeared to move from right side to left side, subjects were instructed to press the corresponding left-sided key (‘Z’ or ‘A’), and when the visual grating appeared to move from left to right, subjects were instructed to press the corresponding right-sided key (‘L’ or ‘M’). If participants correctly responded to the direction of the stimulus, they received positive visual feedback (e.g., images of popular video game cartoon characters were displayed on the computer screen). If participants incorrectly responded to the direction of the stimulus, they received negative visual feedback (e.g., a large red ‘X’ was displayed). Visual gratings appeared on screen for 4 seconds, during which participants could respond. Once the participant responded or at the end of 4 seconds, feedback was presented for 1 second. The following trial would begin 3 second later. All participants completed the first-order visual task, followed immediately by the second-order visual task. For each of the tasks, visual gratings appeared in four blocks of 30 trials, each block consisted of one condition: 180/0, 45/90, 67.5/45, 82.5/15 degrees. The order of the blocks was presented randomly, but participants always received first-order prior to second-order. Prior to administration of the task, participants completed two practice blocks. During the first practice block, a smiley face emoji moved from left to right on the screen (or right to left), and participants were instructed to press the corresponding key based on the direction the smiley faced moved. In the second block, visual gratings at 50/80 angle were presented, and participants pressed key corresponding to direction of movement. Twelve trials of each practice block were administered. If participants did not reach ≥50% correct trials, the block was repeated one time for a total of 24 trials per block. All participants met practice criterion. Depending on the stimulus presented, the subject’s behavioral response was characterized as “Right (similar to Hit)”, “NR (no response)” or “Wrong (similar to FA)”. Since this was a forced two-choice visual discrimination task, a modified d’ (discriminability index) was calculated as follows:

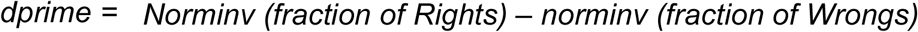

### Statistical analyses

Statistical analyses used are mentioned in the text. For analysis consisting of comparisons between 2 groups a Students t-test (unpaired or paired) was used. For comparisons exceeding 2 groups, a repeated measures ANOVA was performed, followed by the appropriate post hoc tests. Significance was set at p < 0.05. In all the figures, we plot the standard error of the mean (s.e.m.). Graphs either show individual data points from each animal or group means (averaged over different mice) superimposed on individual data points.

## Acknowledgements

The authors thank Kaela Cohan, Steve Cohen and Michael Hong for help with early behavioral experiments, Peyman Golshani and Michael Einstein for advice on mouse behavior, the Janelia GENIE project (GCaMP6s), Peter Yu for building custom lickports, as well as Dean Buonomano, Alcino Silva, Anis Contractor and John Sweeney for feedback on the manuscript. Kim Battista created the illustration in Fig. 1b. This work was supported by the following grants: W81XWH-14-1-0433 (USAMRMC, DOD), Developmental Disabilities Translational Research Program #20160969 (John Merck), SFARI Award 295438 (Simons Foundation) and 5R01HD054453 (NICHD/NIH) awarded to CP-C; K23 MH112936 (NIMH/NIH) to EP; a grant from the Fragile X Alliance of Ohio to CAE; U01 DD001185 (NCBDDD/NIH), U54 HD082008 (NICHD/NIH) and a grant from the Cincinnati Children’s Hospital Research Foundation to EP and CAE.

## Author contributions

A.G. and C.P-C. conceived the project and designed the experiments with help from J.G., L.S. and C.E. for the human studies. A.G. developed the behavioral paradigm for mice and humans. A.G. and D.A.C. wrote the MATLAB code for analysis. A.G, G.C., A.N., L.S. and J.G. conducted the experiments and analyzed the data. A.G., C.E. and C.P-C. interpreted the data and wrote the paper with input from other authors.

## Author information

The authors declare no competing financial interests. Correspondence and requests for materials should be addressed to C.P.-C.

## SUPPLEMENTAL DATA

**Supplemental Figure 1:**
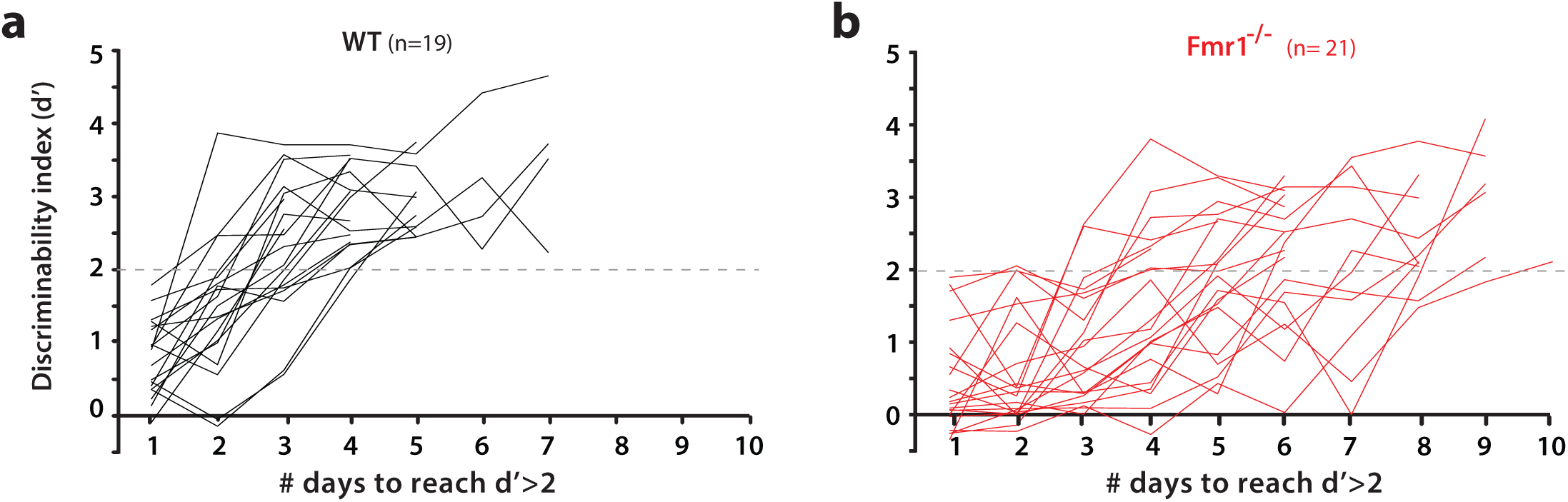
**Behavioral performance on visual discrimination task according to genotype.** (corresponds to data in Fig. 1e-f) a, b. Discriminability index for WT (a, black) and *Fmr1^-/-^* mice (b, red). Each line represents a different mouse. The horizontal dashed line indicates a d’ = 2, which corresponds to 90% correct rejections.

**Supplemental Figure 2:**
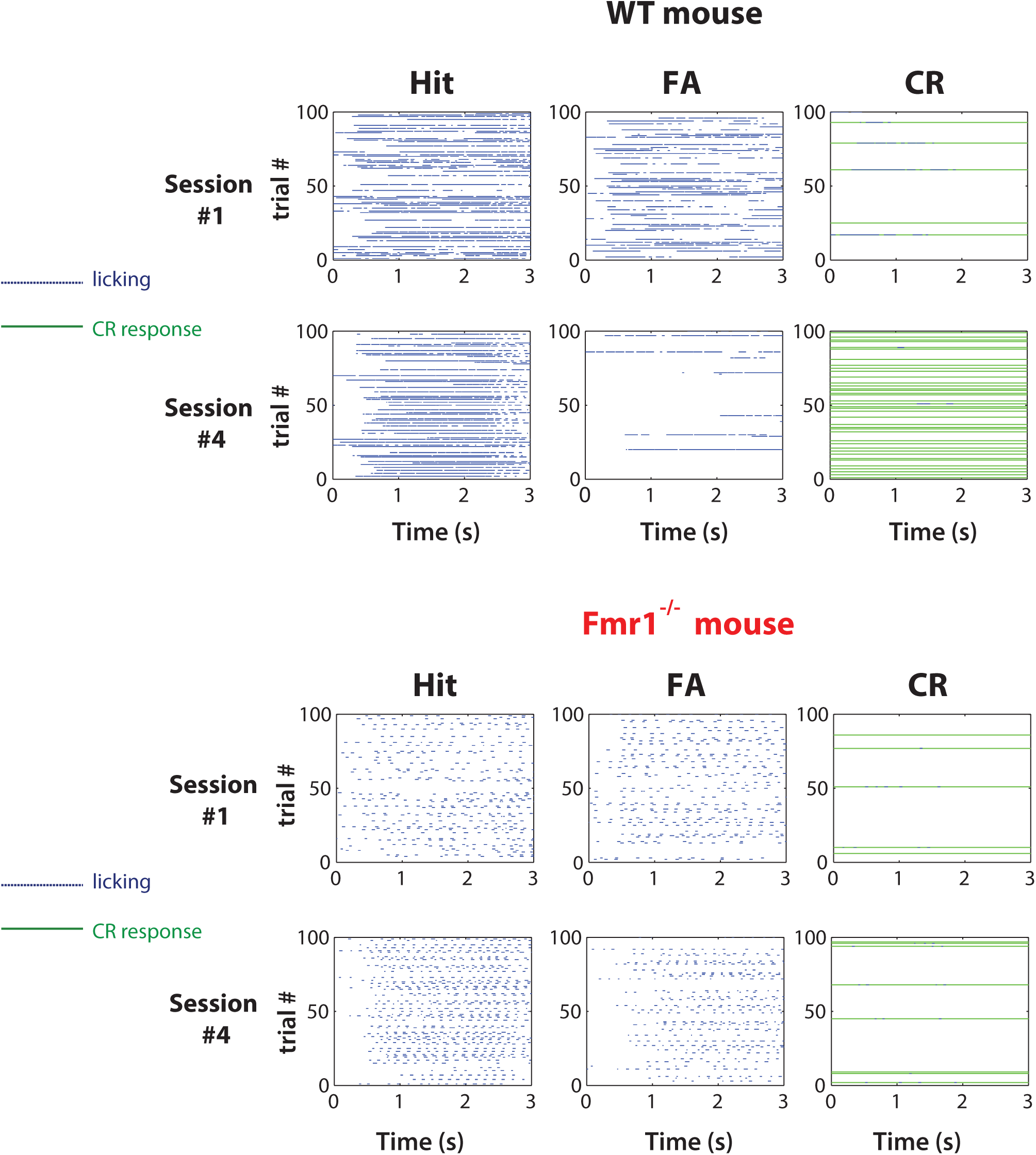
***Fmr1^-/-^* mice show delay in suppressing the No-Go response** (corresponds to data in Fig. 1f) Sample behavioral data for a representative WT mouse (top) and *Fmr1^-/-^* mouse (bottom) over 100 trials in session #1 vs. Session #4. Only ‘Hit’, ‘false alarm’ (FA) and ‘correct rejection (CR) responses are shown. Note how the WT animal is able to suppress the FA responses and increase the proportion of CR responses by session #4, whereas there is virtually no change in response characteristics for the *Fmr1^-/-^* animal between session #1 and session #4.

**Supplemental Figure 3:**
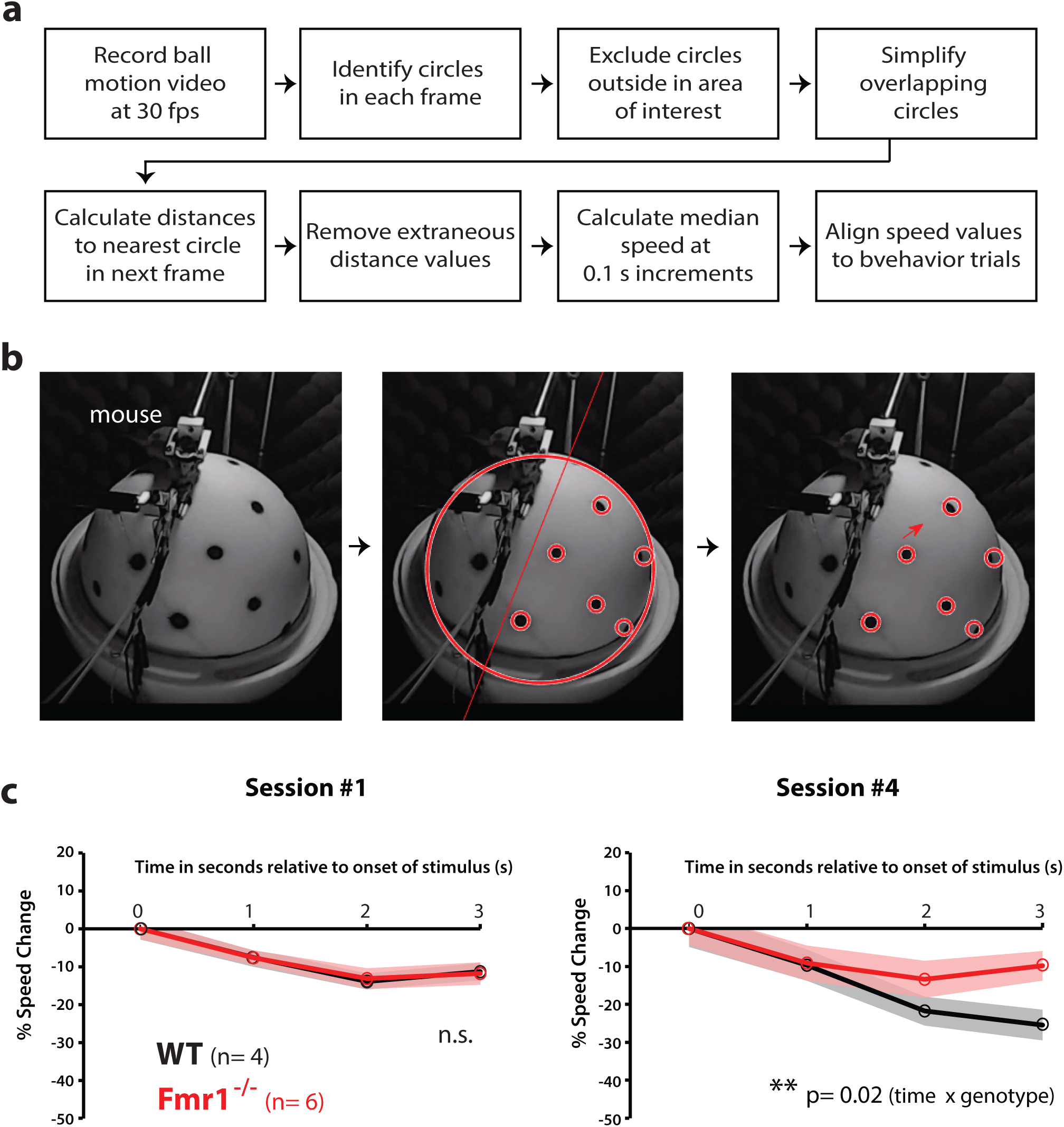
***Fmr1^-/-^* mice do not slow down as much as WT mice during preferred stimuli (Hit responses)** (corresponds to data Fig. 1e, f) a. Workflow for ball motion analysis
b. Analysis of ball motion was done with semi-automated, custom MATLAB scripts to detect black dots painted evenly onto the polystyrene ball.
c. WT and *Fmr1^-/-^* mice showed no obvious differences in running on the ball treadmill during the first session, as determined by ball motion analysis. We observed a significant difference in the degree to which WT and *Fmr1^-/-^* mice slow down during *Hit + Miss* trials (corresponding to preferred stimuli). Data shown represent the average and s.e.m. for n= 4 WT mice and 6 *Fmr1^-/-^* mice.

**Supplemental Figure 4:**
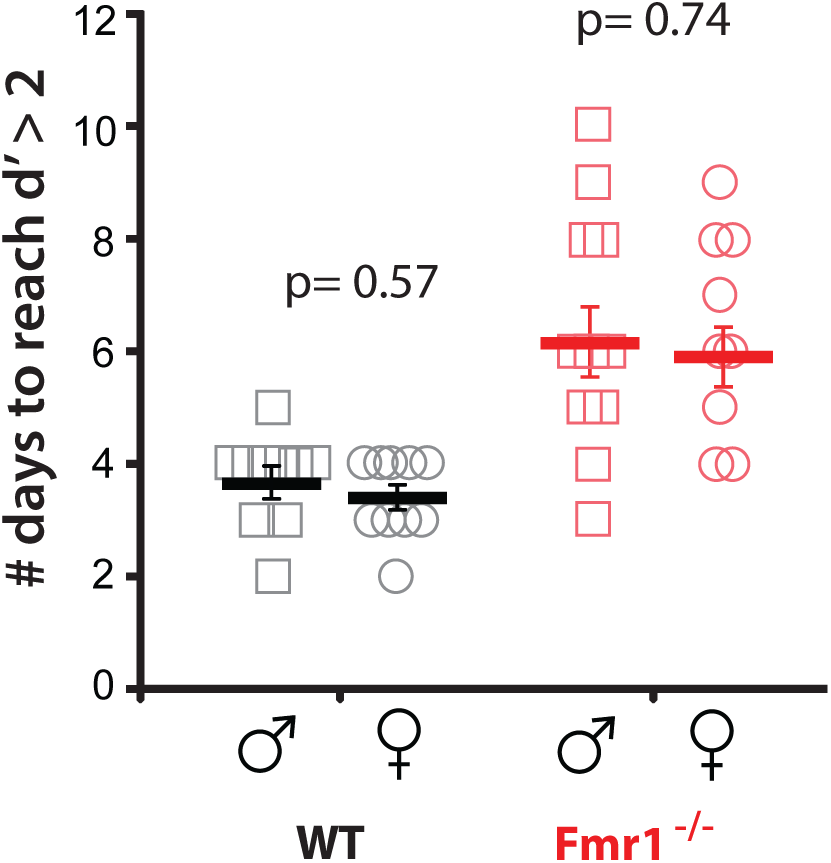
**No sex differences in task performance** (corresponds to data in Fig. 1e-f) There were no significant differences in task performance between male and female mice of either genotype.

**Supplemental Figure 5:**
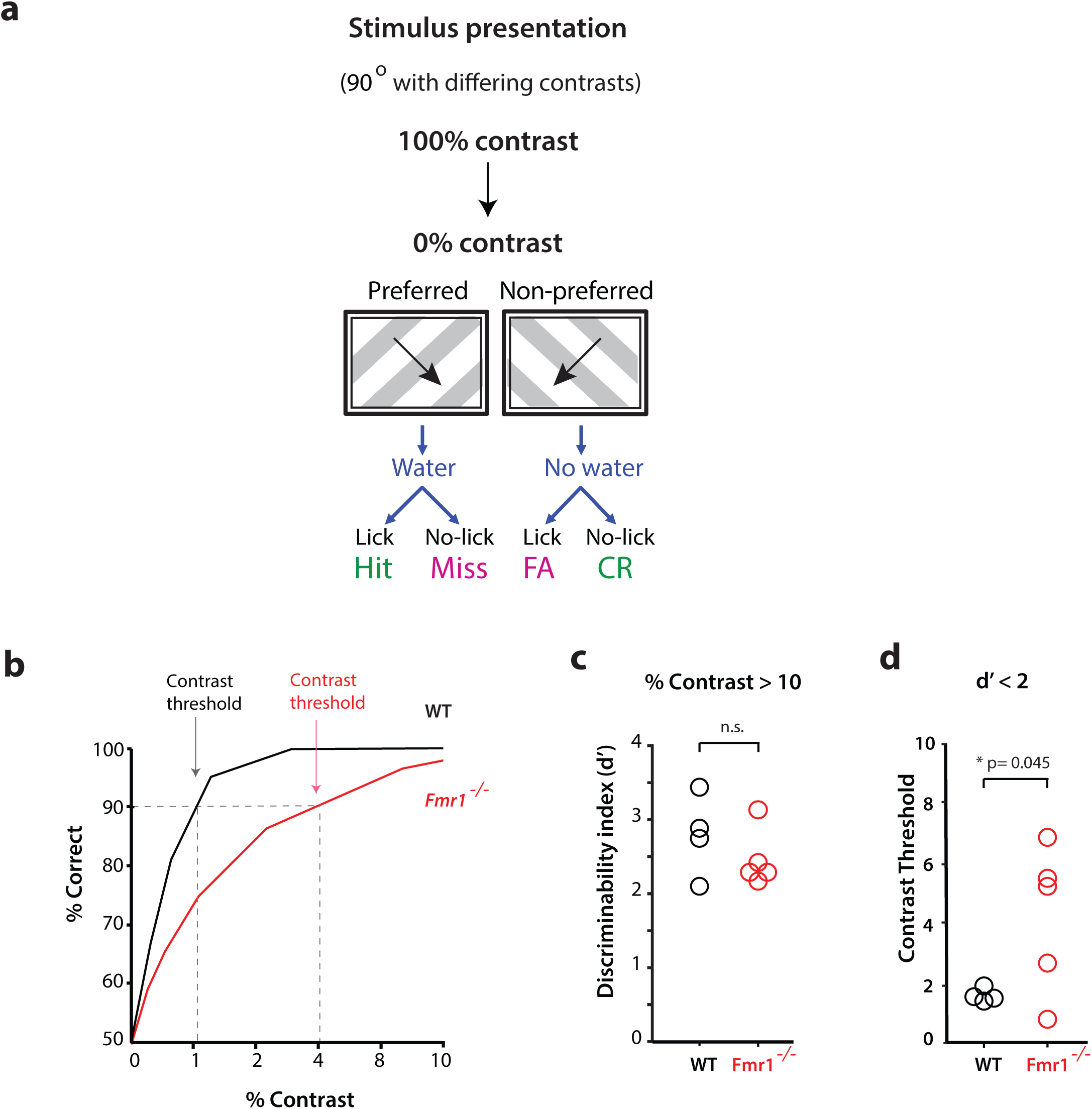
**No deficits in task performance above >10% contrast** (corresponds to data in Fig. 1) a. To investigate whether *Fmr1^-/-^* mice have impairments in basic visual perception, we randomly presented drifting gratings of varying contrast, from 0% to 100% and tested mice of both genotypes on the visual discrimination task with 90^o^ as the angle difference between preferred and non-preferred directions.
b. Representative ** graphs from a WT and a *Fmr1^-/-^* mouse at different contrasts.
c. There were no significant differences in task performance between WT and *Fmr1^-/-^* mice when the contrast for gratings was >10%.
d. However, *Fmr1^-/-^* mice showed a significantly reduced contrast threshold than WT mice to perform at d’>2.

**Supplemental Figure 6:**
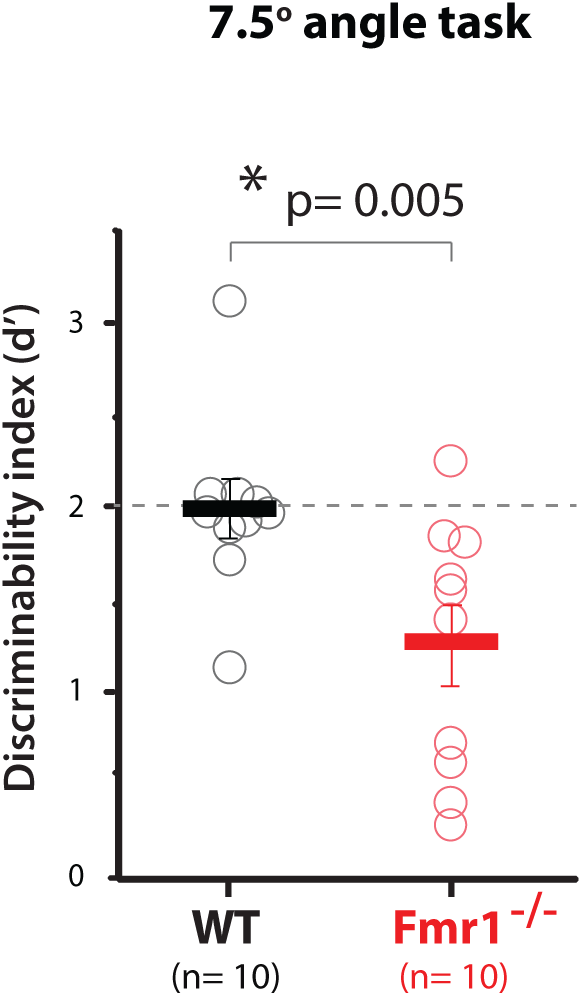
**Reduced angle task at 7.5^o^** (corresponds to data in Fig. 1i) *Fmr1^-/-^* mice show a significantly lower d’ than WT mice when the angle between preferred and non-preferred directions is reduced to 7.5^o^.

**Supplemental Figure 7:**
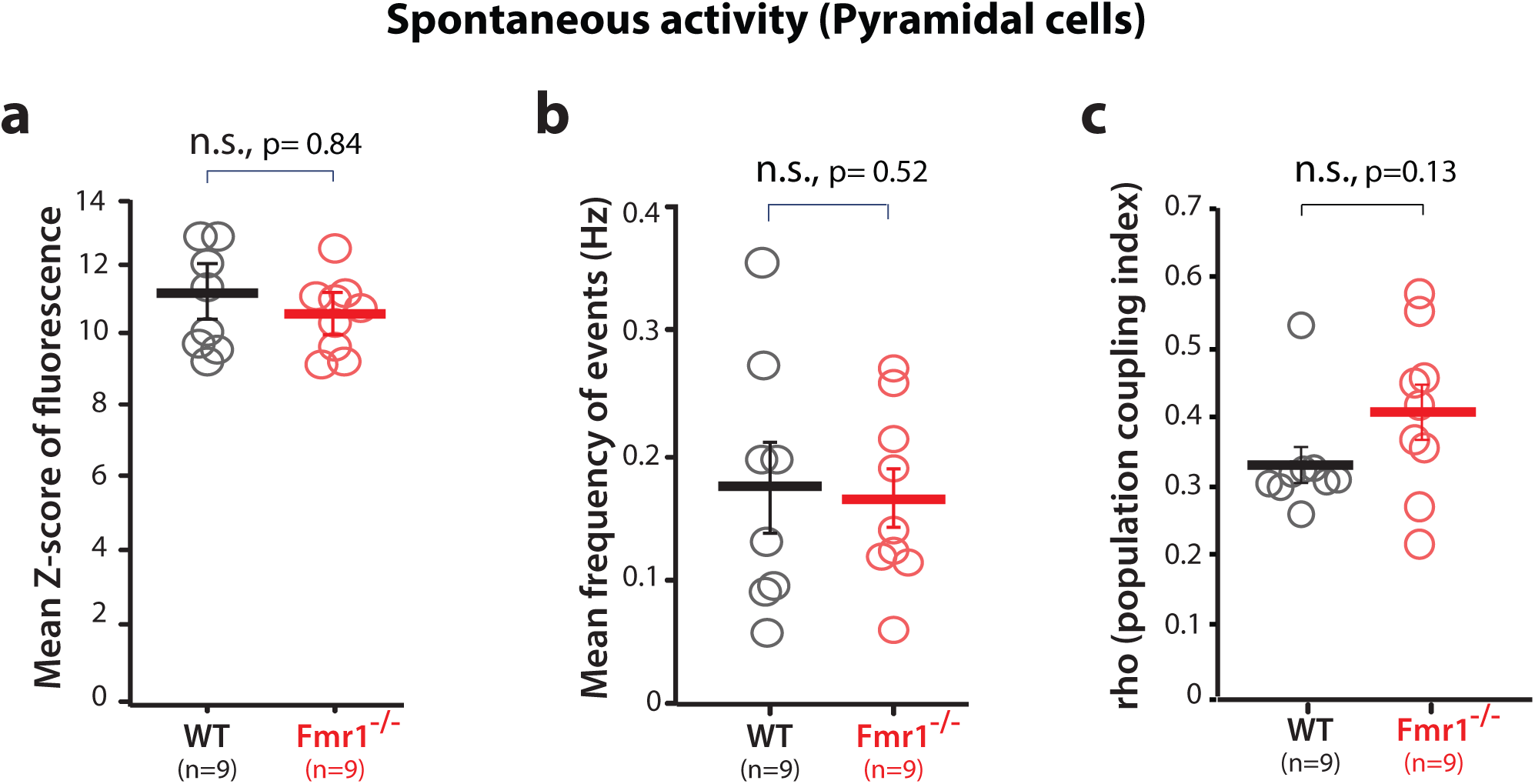
**Calcium imaging of spontaneous activity in pyramidal neurons* in V1** (corresponds to data in Fig. 2) a. There was no significant difference in the amplitude of the calcium signals (as assessed with the mean Z-score of fluorescence) of pyramidal neurons in L2/3 of V1 cortex between WT and *Fmr1^-/-^* mice during recordings of spontaneous activity.
b. There was no significant difference in the frequency of peaks of calcium fluorescence of pyramidal neurons in L2/3 of V1 cortex between WT and *Fmr1^-/-^* mice during recordings of spontaneous activity.
c. There was a non-significant trend toward a higher value for population coupling for pyramidal neurons in L2/3 of V1 cortex in *Fmr1^-/-^* mice compared to WT mice, during recordings of spontaneous activity. * In this legend, the term “pyramidal” refers to all cells recorded using GCaMP6s, the vast majority of which are pyramidal neurons.

**Supplemental Figure 8:**
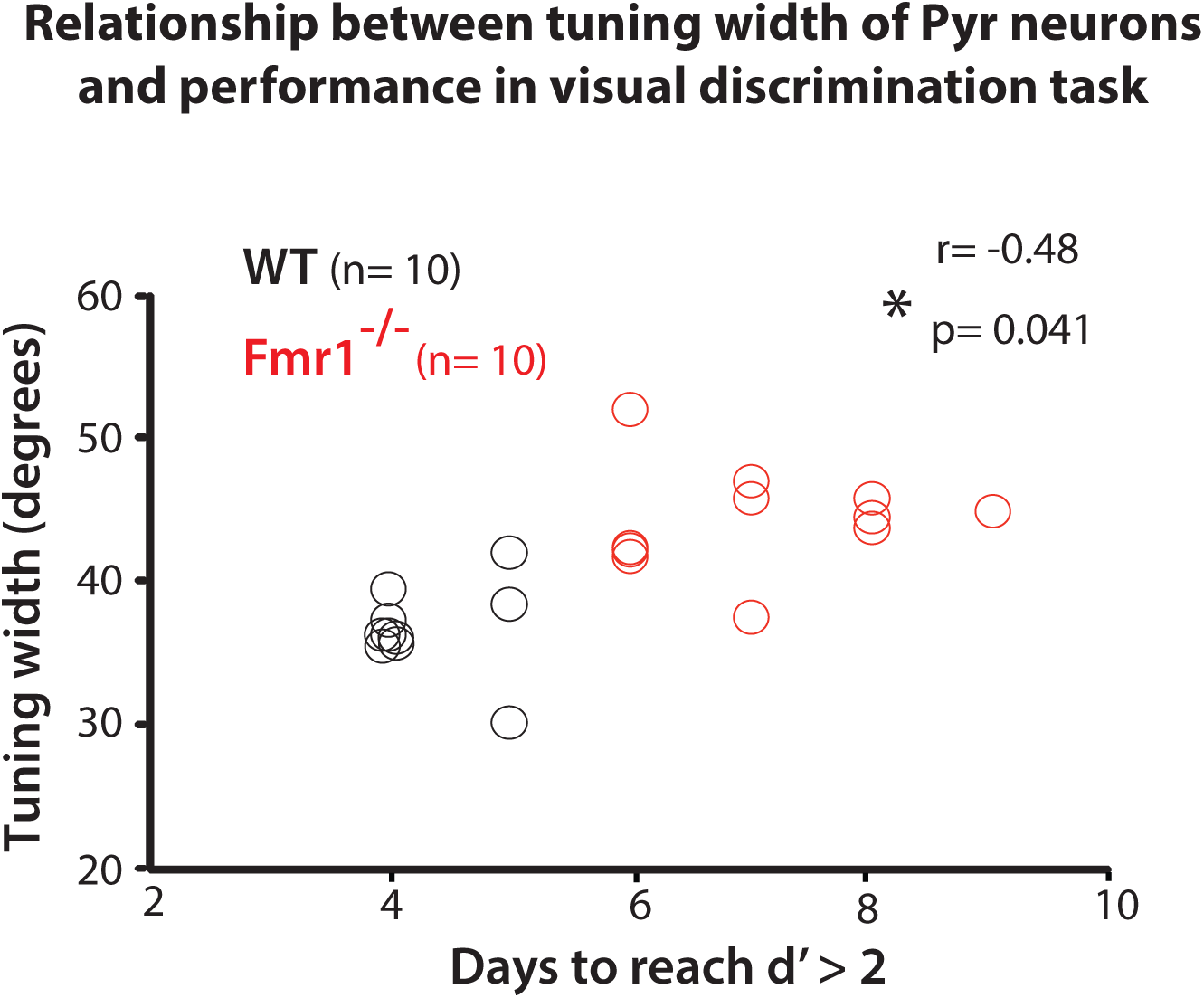
**The tuning width of pyramidal neurons correlates with task performance** (corresponds to data in Fig. 2h) The performance of mice on the visual discrimination task correlates with the tuning width of pyramidal neurons (p= 0.041)

**Supplemental Figure 9:**
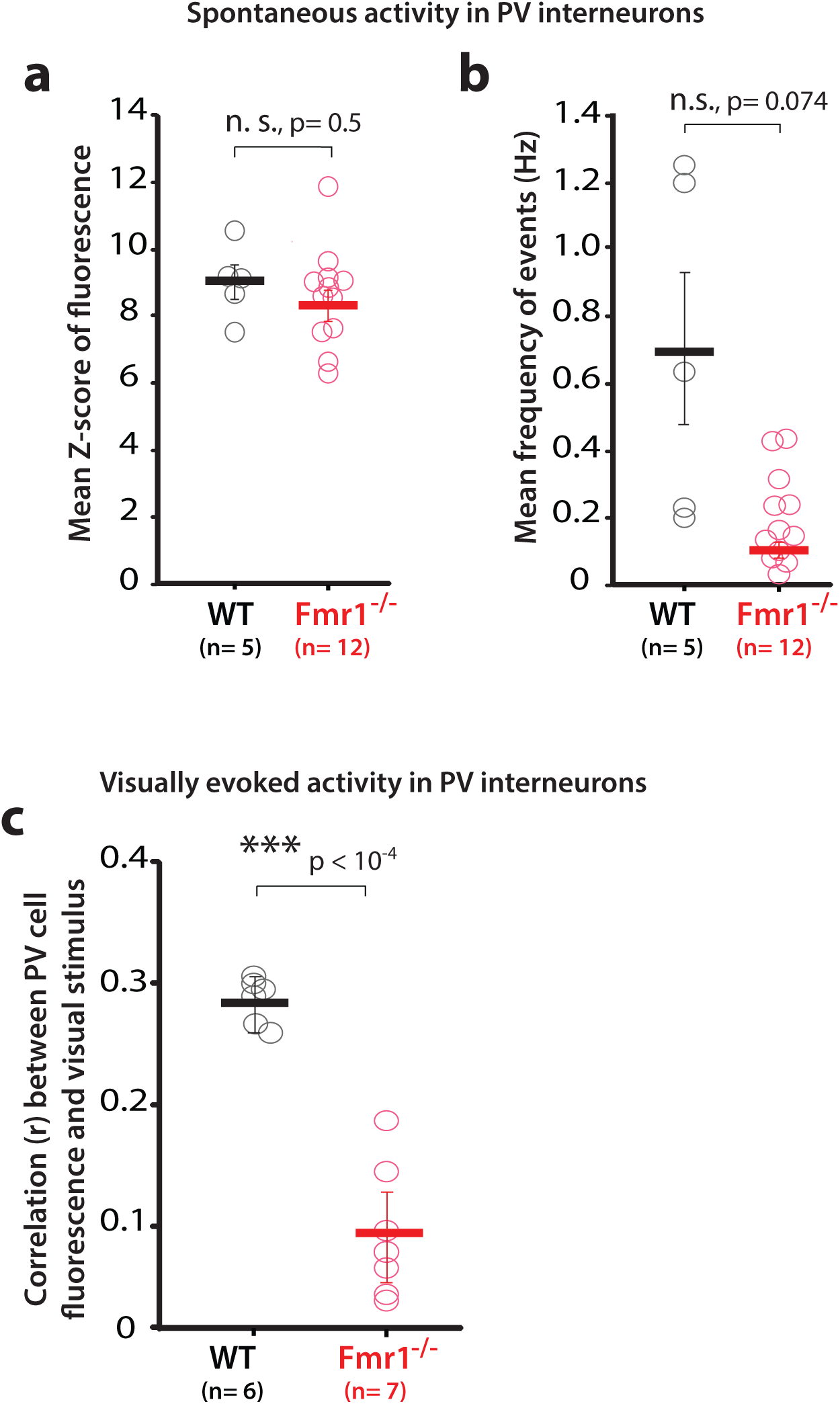
**Calcium imaging of spontaneous activity in PV neurons in V1** (corresponds to data in Fig. 3a-g) a. There was no significant difference in the amplitude of the calcium signals (as assessed with the mean Z-score of fluorescence) of PV neurons in L2/3 between WT and *Fmr1^-/-^* mice during recordings of spontaneous activity in V1.
b. There was a non-significant trend for lower frequency of peaks of calcium fluorescence of PV neurons in L2/3 of *Fmr1^-/-^* mice compared to WT mice, during recordings of spontaneous activity in V1.
c. *Fmr1^-/-^ mice* showed a significantly lower correlation between the activity of PV cells (as assessed by their fluorescence calcium signals) and the epochs of visual stimulation, compared to WT mice.

**Supplemental Figure 10:**
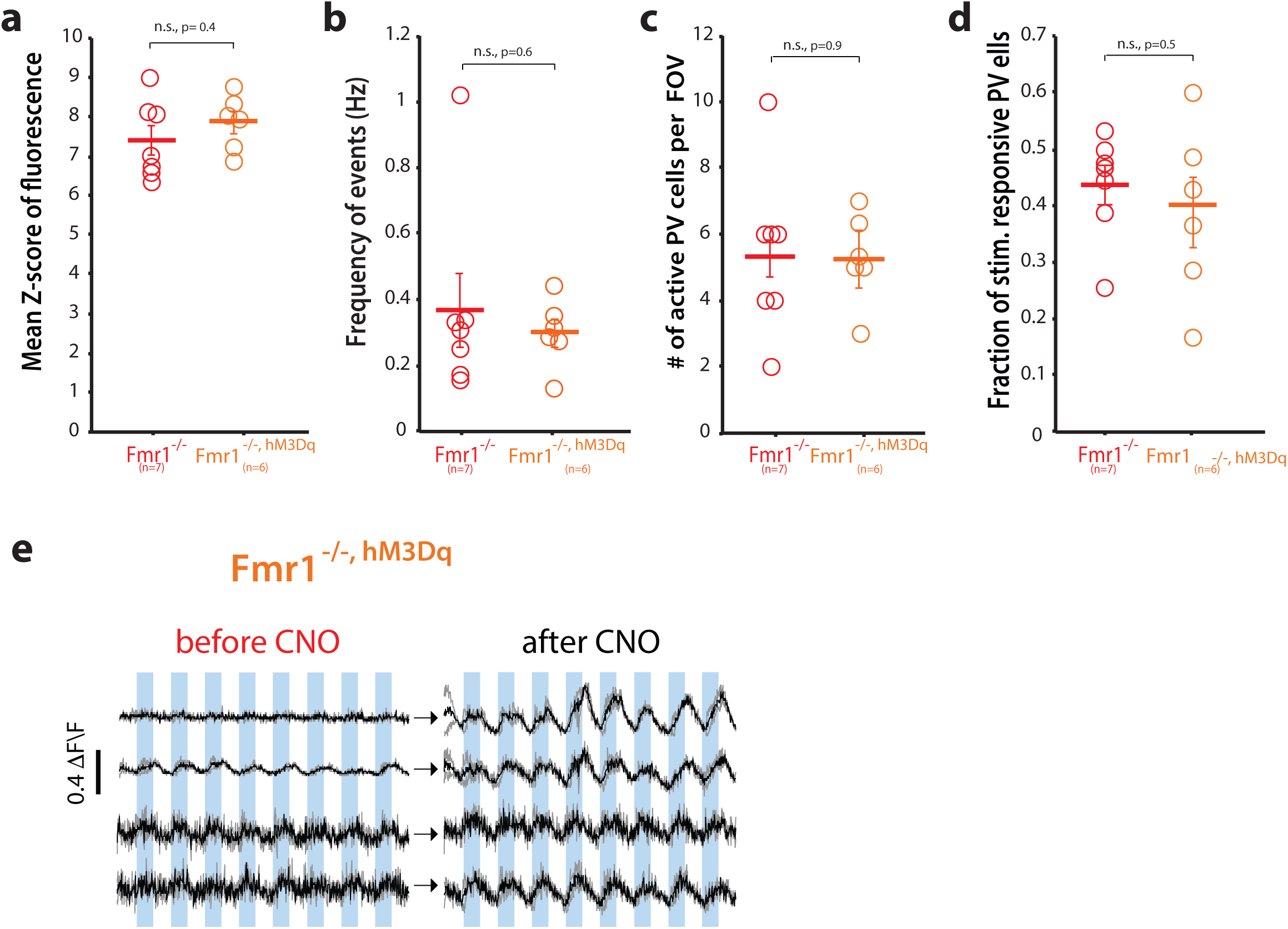
**Controls for DREADD experiments** (corresponds to data in Fig. 3d-g and 3j-l) a-d. DREADD expression does not alter activity of PV cells, as we observed no differences in the amplitude (*a*) or frequency (*b*) of calcium transients, or in the number of PV cells (*d*), or the fraction of PV cells that responded to visual stimuli (i) between *Fmr1^-/-^* mice and *Fmr1^-/-, hM3Dq^* mice (before CNO administration).
e. Example GCaMP6s traces for 4 representative PV neurons in V1 from 4 different *Fmr1^-/-, hM3Dq^* mice before and ~30 min after i.p. injection of clozapine N-oxide (CNO).

**Supplemental Figure 11:**
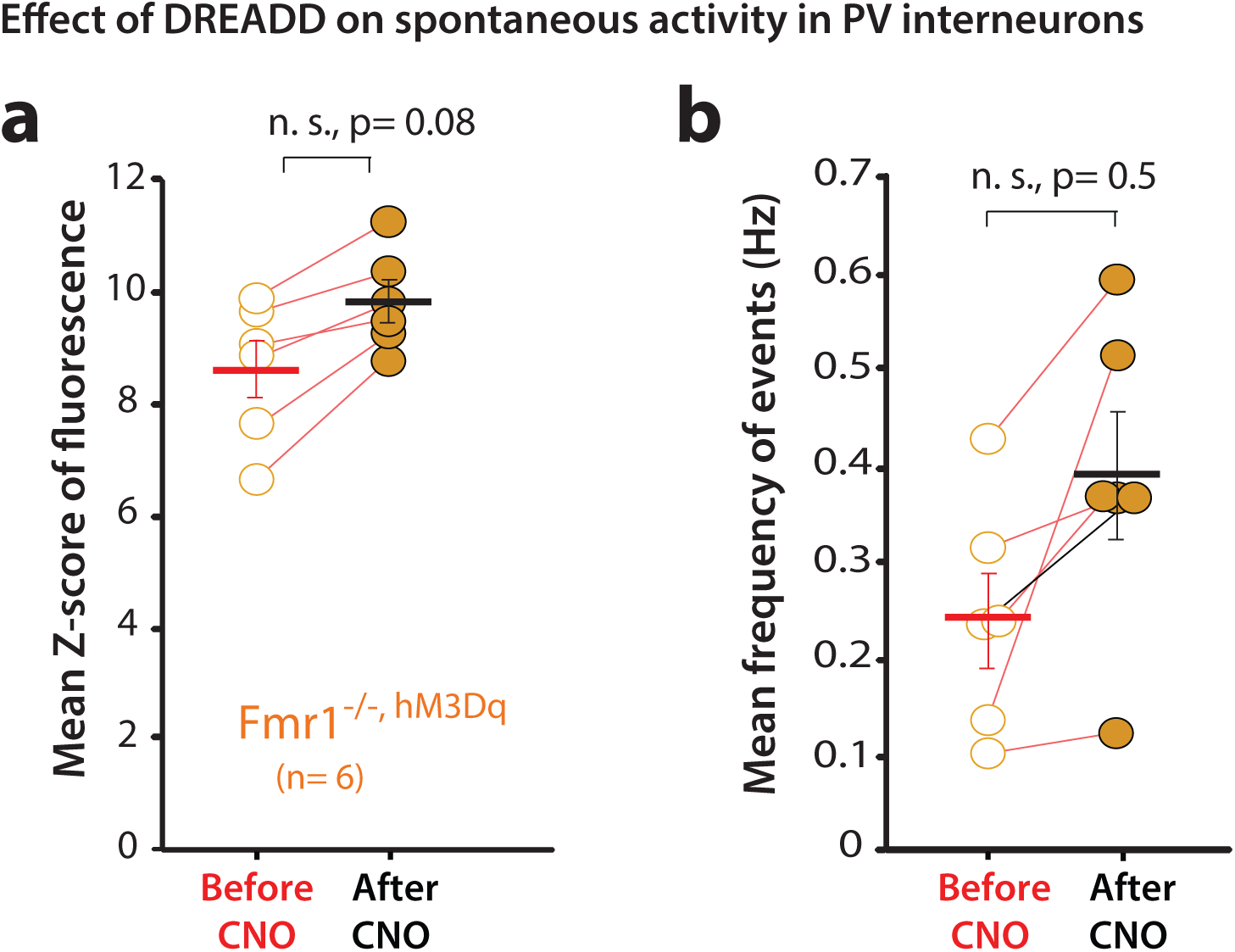
**hM3Dq and CNO effect on spontaneous PV cell activity** (corresponds to data in Fig. 3h-o) a, b. In PV-Cre x *Fmr1^-/-^* mice injected with the hM3Dq DREADD virus in V1, CNO injection led to increases in both the amplitude and the frequency of calcium signals within PV cells, though these changes did not reach significance.

**Supplemental Table 1.** Demographic and clinical information for FXS and control participants.

|  | **FXS**<br>**(n = 8)** | **Control**<br>**(n = 8)** |
| --- | --- | --- |
| **Age in years**<br>*Range* | 30.1 (10.3)<br>15-40 | 29.2 (9.6)<br>15-40 |
| **Abbreviated IQ**<br>*Range* | 52.3 (5.8)<br>47-61 |  |
| **SCQ**<br>*Range* | 12.8 (4.8)<br>7-18 |  |
| **Sensory Profile:**<br>**Seeking/Seeker**<br>*Range* | 25.1 (12.3)<br>3-41 |  |
| **Sensory Profile:**<br>**Avoiding/Avoider**<br>*Range* | 44.8 (20.0)<br>20-71 |  |
| **Sensory Profile:**<br>**Sensitivity/Sensor**<br>*Range* | 40.4 (20.9)<br>11-68 |  |
| **Sensory Profile:**<br>**Registration/Bystander**<br>*Range* | 38.3 (21.7)<br>9-74 |  |
| **Sensory Profile:**<br>**Auditory Processing**<br>*Range* | 19.7 (9.2)<br>6-36 |  |
| **Sensory Profile:**<br>**Visual Processing**<br>*Range* | 11.4 (6.9)<br>1-20 |  |
| **Sensory Profile:**<br>**Touch Processing**<br>*Range* | 16.1 (11.8)<br>2-35 |  |
| **Sensory Profile:**<br>**Movement Processing**<br>*Range* | 8.8 (3.2)<br>2-13 |  |
| **Sensory Profile:**<br>**Body Position Processing**<br>*Range* | 15.0 (12.0)<br>1-35 |  |
| **Sensory Profile:**<br>**Oral Sensory Processing**<br>*Range* | 12.1 (9.8)<br>0-30 |  |

Mean (standard deviation)
IQ = intelligence quotient

**Supplemental Table 2.** Medication type by FXS participant.

| Participant/<br>Drug class | AA | AD | SH | ST |
| --- | --- | --- | --- | --- |
| FXS 1 |  |  | x |  |
| FXS 2 |  |  |  |  |
| FXS 3 |  | x |  |  |
| FXS 4 | x | x |  |  |
| FXS 5 | x | x |  | x |
| FXS 6 | x | x | x |  |
| FXS 7 |  | x |  | x |
| FXS 8 |  |  |  | x |

'x' indicates the participant was taking medication within 48 hours of testing, empty cell indicates no medication
AA: atypical antipsychotic; AD: antidepressant (all SSRI, with exception of FXS 6 who was taking tricyclic); AN: anxiolytic; ST: stimulant

